# A new concept on anti-SARS-CoV-2 vaccines: strong CD8^+^ T-cell immune response in both spleen and lung induced in mice by endogenously engineered extracellular vesicles

**DOI:** 10.1101/2020.12.18.423420

**Authors:** Flavia Ferrantelli, Chiara Chiozzini, Francesco Manfredi, Patrizia Leone, Maurizio Federico

## Abstract

Severe acute respiratory syndrome coronavirus (SARS-CoV)-2 is spreading rapidly in the absence of validated tools to control the growing epidemic besides social distancing and masks. Many efforts are ongoing for the development of vaccines against SARS-CoV-2 since there is an imminent need to develop effective interventions for controlling and preventing SARS-CoV-2 spread. Essentially all vaccines in most advanced phases are based on the induction of antibody response against either whole or part of spike (S) protein. Differently, we developed an original strategy to induce CD8^+^ T cytotoxic lymphocyte (CTL) immunity based on *in vivo* engineering of extracellular vesicles (EVs). We exploited this technology with the aim to identify a clinical candidate defined as DNA vectors expressing SARS-CoV-2 antigens inducing a robust CD8^+^ T-cell response. This is a new vaccination approach employing a DNA expression vector encoding a biologically inactive HIV-1 Nef protein (Nef^mut^) showing an unusually high efficiency of incorporation into EVs even when foreign polypeptides are fused to its C-terminus. Nanovesicles containing Nef^mut^-fused antigens released by muscle cells are internalized by antigen-presenting cells leading to cross-presentation of the associated antigens thereby priming of antigen-specific CD8^+^ T-cells. To apply this technology to a design of anti-SARS-CoV-2 vaccine, we recovered DNA vectors expressing the products of fusion between Nef^mut^ and four viral antigens, namely N- and C-terminal moieties of S (referred to as S1 and S2), M, and N. All fusion products are efficiently uploaded in EVs. When the respective DNA vectors were injected in mice, a strong antigen-specific CD8^+^ T cell immunity was generated. Most important, high levels of virus-specific CD8^+^ T cells were found in bronchoalveolar lavages of immunized mice. Co-injection of DNA vectors expressing the diverse SARS-CoV-2 antigens resulted in additive immune responses in both spleen and lung. EVs engineered with SARS-CoV-2 antigens proved immunogenic also in the human system through cross-priming assays carried out with *ex vivo* human cells. Hence, DNA vectors expressing Nef^mut^-based fusion proteins can be proposed as anti-SARS-CoV-2 vaccine candidates.

## Introduction

Severe acute respiratory syndrome coronavirus (SARS-CoV)-2 first emerged in late 2019 in China (*1–3*). Globally, the virus has since infected over 72 million individuals and caused more than 1.6 million deaths (*4*). Given the severity of the disease, vaccines and therapeutics to tackle this novel virus are urgently needed.

The majority of SARS-CoV-2 vaccines currently in development aims at inducing neutralizing antibodies against either whole or part of virus spike (S) protein. However, follow-up studies from patients who recovered from previous epidemic of SARS-CoV suggest that specific antibody responses are short lived with few or no memory B-cells (*5–12*), and target the primary homologous strain. Conversely, memory T cells against SARS-CoV persist 11 years post-infection and have the potential to induce cross-reactive immunity (*13–17*). An increasing number of studies shows that also SARS-CoV-2 convalescent patients develop robust T-cell responses (*18–22*). Besides S protein, SARS-CoV-2 membrane (M) protein also elicits strong CD8^+^ T cell responses, and a significant reactivity was also reported for both nucleocapsid (N) (*18*) and ORF1ab (*19*) proteins. Taken together, these evidences suggest that a long-lasting, broad vaccine formulation against SARS-CoV-2 could and should induce strong memory T cell responses against multiple viral antigens.

Extracellular vesicles (EVs) are a heterogeneous population of membrane nanovesicles among which exosomes are the most studied (*23*). EVs are released by most cell types including myocytes (*24*), and have critical roles in cell-to-cell communication and regulation of immune responses (*25–28*).

We previously described the ability of Human Immunodeficiency virus (HIV)-1 Nef to incorporate into EVs released by multiple cell types including CD4^+^ T lymphocytes and dendritic cells (*29*). The incorporation into EVs increases by approximately 100-fold in the case of the Nef G3C-V153L-E177G mutant, most likely due to increased stabilization with cell membranes (*30, 31*). Moreover, and extremely beneficial for a vaccine, V153L-E177G mutations make Nef^mut^ defective for basically all detrimental Nef functions including both CD4 and MHC Class I down-regulation, increased HIV-1 infectivity, and p21 activated kinases (PAK)-2 activation (*31, 32*). Furthermore, we observed that the efficiency of Nef^mut^ incorporation into EVs is maintained even when a foreign protein is fused to its C-terminus (*30, 31, 33–36*). When DNA vectors expressing Nef^mut^-based fusion proteins are intramuscularly (i.m.) injected in mice, detectable amounts of the fusion protein are packed into EVs while not altering their spontaneous release from muscle tissue. Nef^mut^-fused antigens released inside muscle derived-EVs are then internalized by antigen-presenting cells (APCs) which, in turn, cross-present EV content to activate antigen-specific T cells. These *in vivo* engineered EVs are assumed to freely circulate into the body, thereby acting as an effective vaccine by eliciting potent antigen-specific CTL responses (*30, 31, 36)*. The effectiveness and flexibility of this vaccine platform has been demonstrated with an array of viral products of various origins and sizes, including but not limited to: Human Papilloma Virus (HPV)16-E6 and −E7; Ebola Virus VP24, VP40 and NP; Hepatitis C Virus NS3; West Nile Virus NS3; and Crimean-Congo Hemorrhagic Fever NP. Of note, in our hands very low to undetectable antigen-specific CD8^+^ T immune responses were observed when animals were injected with DNA vectors expressing the antigen open reading frame (ORF) devoid of the Nef^mut^ sequences (*30, 36*). On the other hand, no antibody response was detected against Nef or any of the antigens fused to it in mice injected with Nef^mut^-based DNA vectors.

We tested three SARS-CoV-2 structural antigens, namely spike (S), membrane (M), and nucleocapsid (N) proteins in the context of the Nef^mut^ system. The immunogenicity of DNA vectors expressing each SARS-CoV-2 protein fused with Nef^mut^ and injected in mice either alone or in combination was evaluated in both spleens and lung airways. The immunogenicity of engineered EVs was tested also in *ex vivo* human blood cells.

## Results

### Construction of vectors expressing Nef^mut^ fused with SARS-CoV-2 antigens

All Nef^mut^/SARS-CoV-2 fusion proteins were expressed in the context of the pVAX1 vector (Fig. 1A). The ORFs encoding SARS-CoV-2 S, M and N proteins were from the Italian clinical isolate of SARS-CoV-2 ITA/INMI1/2020 (https://www.ncbi.nlm.nih.gov/nuccore/MT066156; GenBank: MT066156.1). Each Nef^mut^-fusion construct has been designed to guarantee optimal internalization into EVs meanwhile preserving already characterized mouse immunodominant epitopes. SARS-CoV-2 amino acid sequences included in the Nef^mut^-based fusion proteins are highlighted in fig. 1B.

**Figure 1.**
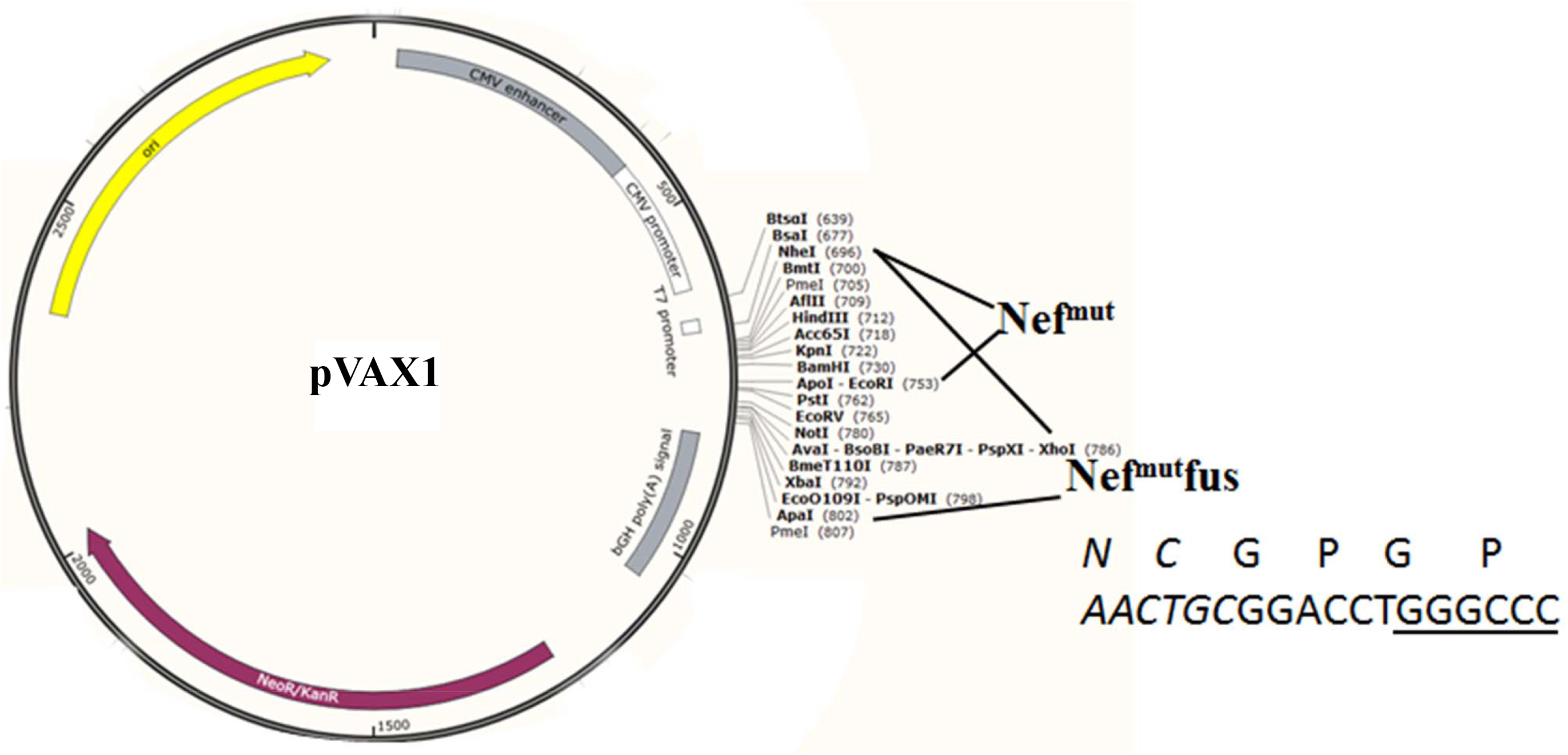
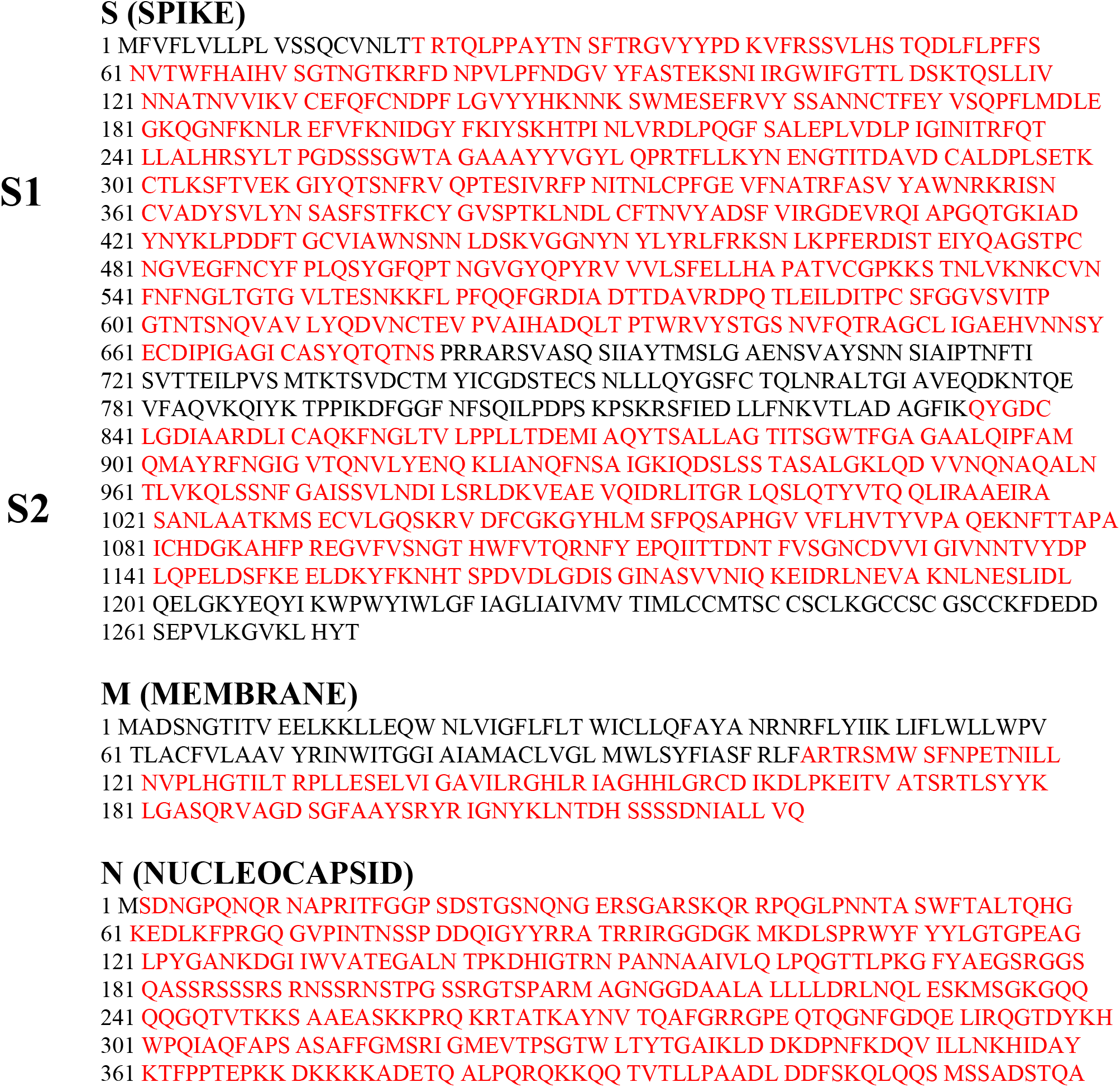
Map of vectors expressing SARS-CoV-2-based fusion proteins, and relative amino acid sequences. A. Shown are the map of pVAX1 vector as well as the cloning strategies pursued to obtain both pVAX1-Nef^mut^ and pVAX1-Nef^mut^fusion vectors. Restriction sites where the SARS-CoV-2 ORFs were inserted are also indicated. On the bottom right, shown are both amino acid and nucleotide sequences of Nef^mut^ C-terminus in the pVAX1-Nef^mut^fusion vector (in italics), as well as of the GPGP linker overlapping the *Apa* I restriction site (underlined sequence). B. Amino acid sequences of S, N, and M proteins from the Italian clinical isolate of SARS-CoV-2 ITA/INMI1/2020 (https://www.ncbi.nlm.nih.gov/nuccore/MT066156; GenBank: MT066156.1). Amino acid sequences included in the Nef^mut^-based vectors are highlighted.

In SARS-CoV-2, S is cleaved at the boundary between S1 and S2 subunits which remain non-covalently bound in the pre-fusion conformation (*37*). The cleavage occurs at the furin-like site PRRARS. We predicted that the furin-dependent cleavage of S would negatively affect the uploading into EVs of the entire S protein fused with Nef^mut^. To overcome this limitation, we designed two Nef^mut^-based constructs, i.e, Nef^mut^-S1 (aa 19 to 680) where both signal peptide and furin-like cleavage site were excluded, and Nef^mut^-S2 (aa 836 to 1196), including the extracellular portion of the protein with the exclusion of the two fusion domains, the transmembrane region, and the short intracytoplasmic tail.

The SARS-CoV-2 M protein (221 aa) is composed of an amino-terminal exterior region of 18 amino acids, a transmembrane region accounting for approximately one third of the entire protein, and a C-terminal region composed of 123 intraluminal residues (*37*). To guarantee an efficient EV uploading, only the C-terminal region of M (aa 94 to 221) was fused to Nef^mut^.

Finally, the full length ORF of the N protein (422 aa), except for M1 amino acid, was fused to Nef^mut^.

### Nef^mut^-based products of fusion with SARS-CoV-2 antigens are efficiently uploaded in EVs

The cell expression of the products of fusion between Nef^mut^ and SARS-CoV-2 antigens S1, S2, M, and N was evaluated by transient transfection in HEK293T cells. To inspect the uploading into EVs of the fusion products, supernatants from transfected cell were collected 48-72 h after transfection, and then processed by differential centrifugations.

Both cell lysates and EVs isolated from the respective supernatants were analyzed by western blot assay (Fig. 2). Upon incubation with anti-Nef Abs, the cell-associated steady-state levels of all Nef^mut^-derivatives were clearly detectable. The strongest signals appeared in lysates of cells expressing either Nef^mut^ alone or its product of fusion with N. In this case, products of lower molecular weight were also detectable, possibly originated by intracellular cleavage.

**Figure 2.**
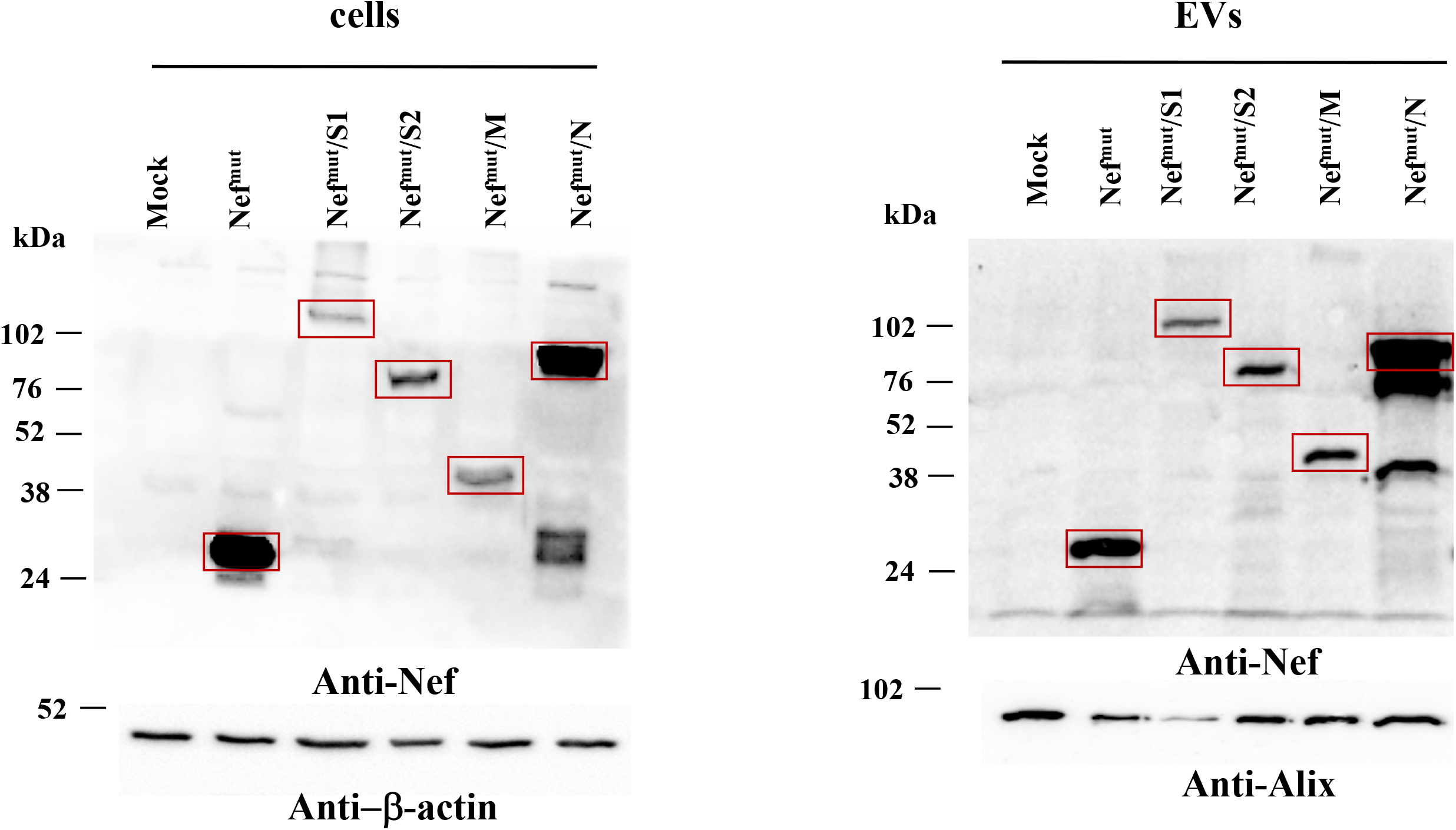
Detection of Nef^mut^/SARS-CoV-2 fusion products in transfected cells and EVs. Western blot analysis on 30 μg of lysates from HEK293T cells transfected with DNA vectors expressing Nef^mut^ fused with either S1, S2, M or N SARS-CoV-2 ORFs (left panels). Equal volumes of buffer where purified EVs were resuspended after differential centrifugations of the respective supernatants were also analyzed (right panels). As control, conditions from mock-transfected cells as well as cells transfected with a vector expressing Nef^mut^ alone were included. Polyclonal anti-Nef Abs served to detect Nef^mut^-based products, while β-actin and Alix were revealed as markers for cell lysates and EVs, respectively. Relevant signals are highlighted. Molecular markers are given in kDa. The results are representative of four independent experiments.

The results we obtained from the analysis of EVs basically reflected those from cell lysate analysis (Fig. 2). Also in this case, the presence of apparently full-length Nef^mut^/N fusion product coupled with that of two products of lower molecular weight.

Taken together, these results indicated that all fusion products we designed are stable and associate with EVs.

### Virus-specific CD8^+^ T cells were detected in spleens from mice injected with vectors expressing Nef^mut^ fused with each SARS-CoV-2 antigen

Next, we evaluated the immunogenicity of DNA vectors expressing each SARS-CoV-2 antigen fused with Nef^mut^. As benchmark of CD8^+^ T cell immunity induced through the Nef^mut^ system, mice were immunized with a vector expressing Nef^mut^/E7, i.e., a vector whose injection generates a both strong and effective anti-E7 CTL immune response (*36, 38*). Either C57 Bl/6 or, in the case of immunization with the Nef^mut^/S2 vector, Balb/c mice were i.m. inoculated in each quadriceps with 10 μg of each DNA vector and, as control, equal amounts of either void or pVAX1-Nef^mut^ vector. Injections were immediately followed by electroporation procedures. The immunizations were repeated 14 days later and, after additional 14 days, mice were sacrificed. Splenocytes were then isolated and cultured overnight in IFN-γ EliSpot microwells in the presence of either unrelated or antigen-specific octo-decamers specific for CD8^+^ T cell immunodominant epitopes already described for SARS-CoV, and whose sequences are unmodified in SARS-CoV-2 antigens. Both sustained and comparable antigen-specific CD8^+^ T cell activation were observed in splenocyte cultures from mice inoculated with each vector expressing the diverse Nef^mut^-based fusion proteins (Fig. 3). Although the assay does not allow a rigorous quantification of the immune response, the CD8^+^ T cell activation extents detected in mice injected with vectors expressing SARS-CoV-2-derivatives appeared comparable to that induced by the Nef^mut^/E7-expressing vector we considered as a “gold-standard”.

**Figure 3.**
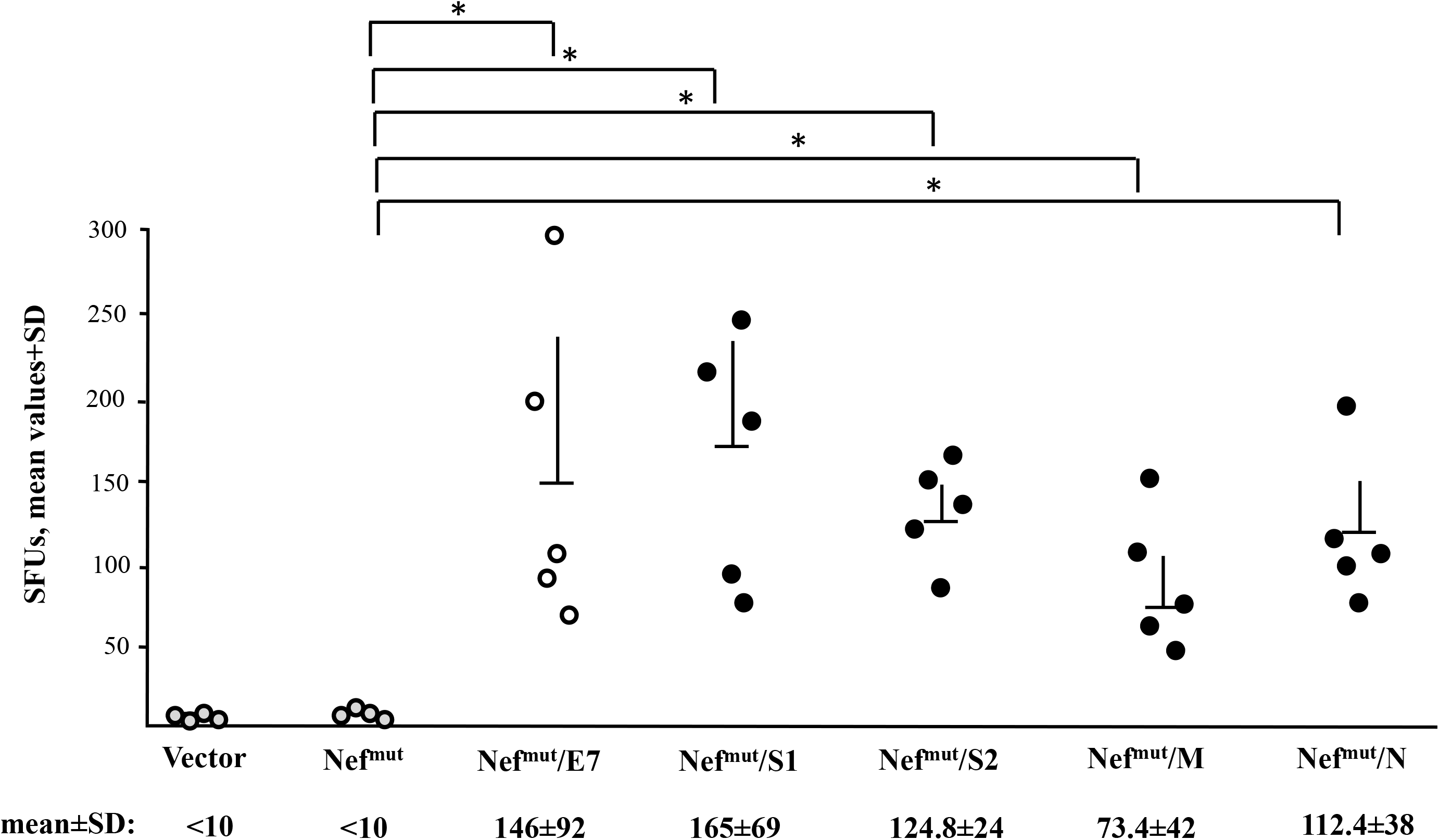
SARS-CoV-2-specific CD8^+^ T cell immunity induced in spleens from mice after i.m. DNA injection. CD8^+^ T cell immune response in either C57 Bl/6 or, for immunization with S2-expressing vector only, Balb/c mice inoculated i.m. with the DNA vectors expressing Nef^mut^ either alone (4 mice) or fused with the indicated SARS-CoV-2 antigens (5 mice per group). As controls, mice were inoculated with either void vector (4 mice) or, for C57 Bl/6 mice only, a vector expressing Nef^mut^/E7 (5 mice). At the time of sacrifice, 2.5×10^5^ splenocytes were incubated o.n. with or without 5 μg/ml of either unrelated or SARS-CoV-2-specific peptides in triplicate IFN-γ EliSpot microwells. Shown are the numbers of IFN-γ spot-forming units (SFU)/well calculated as mean values of triplicates after subtraction of mean spot numbers calculated in wells of splenocytes treated with unspecific peptides. Reported are intragroup mean values + standard deviations also. The SARS-CoV-2-specific immune response in Balb/c mice injected with either void or Nef^mut^ expressing vectors remained at background levels, i.e., the levels measured in wells treated with unspecific peptides. Statistical significance compared to values obtained with splenocytes injected with Nef^mut^ expressing vector was determined by Mann-Whitney U test. \**p*<0.05.

These data indicated that all four SARS-CoV-2 based DNA vectors were able to elicit a virus-specific CD8^+^ T cell immunity.

### Detection of SARS-CoV-2-specific CD8^+^ T lymphocytes in cells from bronchoalveolar lavages of immunized mice

The induction of a CD8^+^ T cell immune response at the pulmonary tissues should be considered a mandatory feature for any CD8^+^ T cell-based vaccine against respiratory diseases. In general, the immune response against respiratory viruses leads to the formation of three distinct antigen-specific CD8^+^ T cell populations: circulating effector memory (T_EM_); central memory (T_CM_), basically residing in secondary self-renewing cell population only minimally replenished by circulating T_EM_ (*40*). On this basis, the presence of virus-specific CD8^+^ T cells in spleens would not necessarily guarantee adequate levels of immunity at lung tissues, i.e., the district directly involved in SARS-CoV-2 pathogenesis.

The here above immunogenicity experiment was reproduced with the specific aim to test the cell immune response at the level of lung airways. To this end, mice were immunized as described and, 14 days after the second immunization, cells from both spleens and bronchoalveolar lavages (BALs) were isolated and tested in IFN-γ EliSpot assay. Either pools of peptides to test the total cell immune response, or octo-decamers specific for the CD8^+^ T cell immune response were used. Results obtained with splenocytes (Fig. 4A) fairly reproduced those described in the previous immunogenicity experiment. The higher responses detected with peptide pools were likely consequence of a co-induced CD4^+^ T cell immune response. Concerning the immune responses detectable in cells isolated from BALs (Fig. 4B), quite high percentages of SARS-CoV-2 specific CD8^+^ T cells compared to the number of activated PMA cells were found (Fig. 4C). Using peptide pools, the immune responses appeared to be increased less significantly compared to what observed with splenocytes (Fig. 4C), likely consequence of the predominance of CD8^+^ T cell immunity. This hypothesis was supported by data obtained through intracellular staining (ICS) and FACS analysis of cells from BALs from mice immunized by Nef^mut^/S1 vector (Fig. 4D). After stimulation with the pool of S1 peptides, a prevalent activation of CD8^+^ T over CD4^+^ T cells was observed. Of note, a significant subpopulation of activated CD8^+^ T cells co-expressed IFN-γ, IL-2, and TNF-α indicating the induction of polyfunctional CD8^+^ T cells (Fig. 4D)..

**Figure 4.**
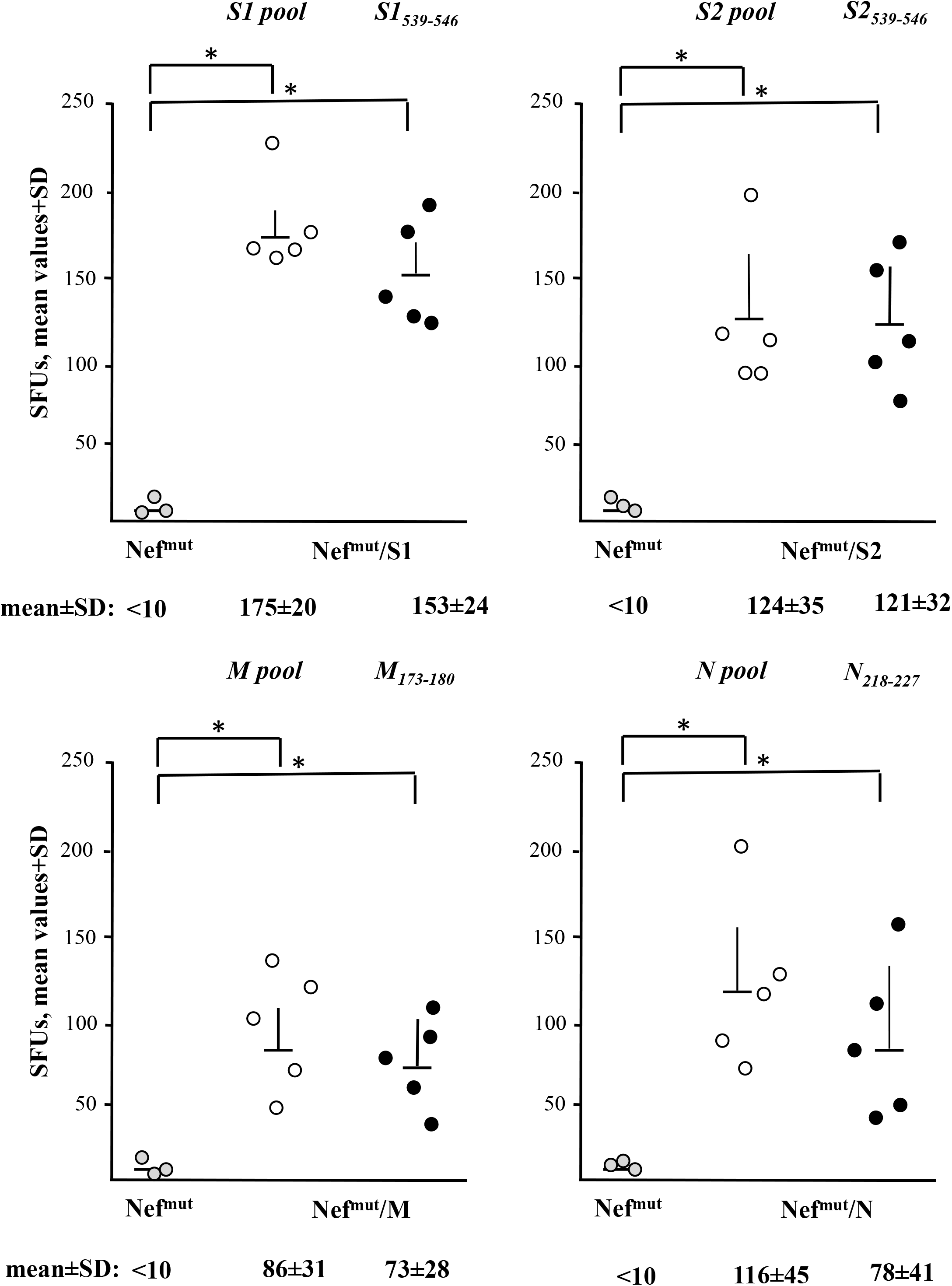
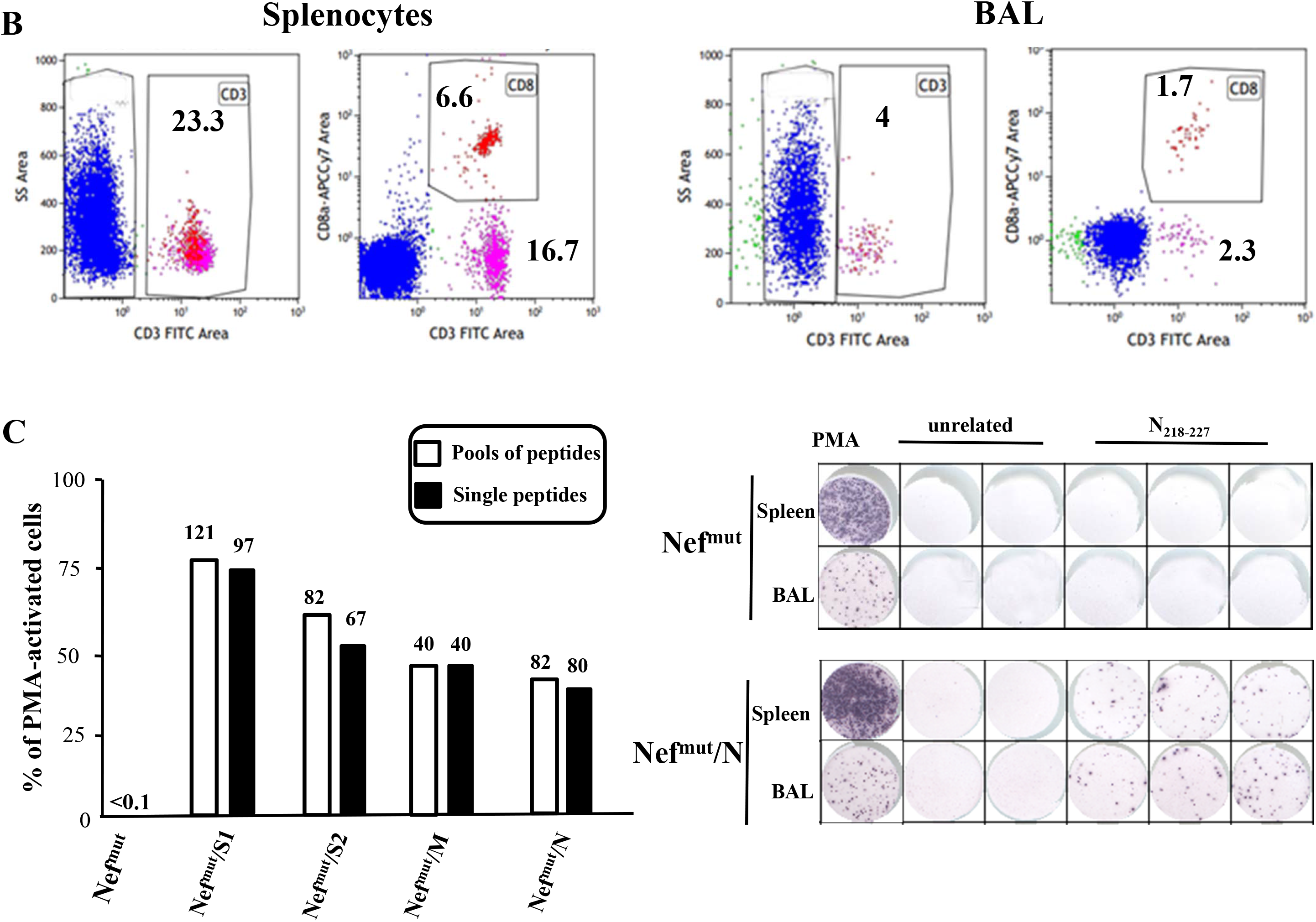
SARS-CoV-2-specific total cell or CD8^+^ T cell immunity in both spleens and BALs of mice after i.m. DNA injection. A. Total cell or CD8^+^ T cell immune responses in C57 Bl/6 and, for S2 immunization only, Balb/c mice inoculated i.m. with the DNA vectors expressing Nef^mut^ either alone (3 mice) or fused with the indicated SARS-CoV-2 antigens (5 mice per group). The total cell immune responses were measured using pools of 13-to 17-mers, whereas single octo-decamers were used to evaluate the CD8^+^ T cell immune responses. At the time of sacrifice, 2.5×10^5^ splenocytes were incubated o.n. with or without 5 μg/ml of either unrelated or SARS-CoV-2-specific peptides in triplicate IFN-γ EliSpot microwells. Shown are the numbers of IFN-γ SFU/well as mean values of triplicates after subtraction of mean spot numbers measured in wells of splenocytes treated with unspecific peptides. Reported are intragroup mean values + standard deviations also. No virus-specific immune responses were detected using a pool of peptides from a heterologous viral product, i.e., HCV-NS3. Statistical significance compared to values obtained with splenocytes injected with Nef^mut^ expressing vector was determined by Mann-Whitney U test. \**p*<0.05. B. FACS analysis of cell populations recovered through BALs (right panels) compared to those from spleens (left panels). Total lymphocytes were identified through anti-CD3 labeling among which both CD4^+^ and CD8^+^ sub-populations were then distinguished. Percentages of CD3^+^, CD4^+^, and CD8^+^ over the total of events are indicated. The data refer to mice injected with the Nef^mut^/N expressing vector, and are representative of eight independent analyses. C. Percentages of SFUs detected in IFN-γ EliSpot microwells seeded with 10^5^ cells from pooled BALs in the presence of virus-specific peptides compared to PMA plus ionomycin. Cells were treated with either pools or single peptides. Cell samples seeded with unrelated peptides scored at background levels. Absolute SFU numbers are also indicated. The results are representative of two independent experim Relevant signals are highlighted. ents. On the right, shown are raw data from a representative IFN-γ EliSpot plate where cells from both spleens and BALs of mice injected with DNA vectors expressing either Nef^mut^ or Nef^mut^/N were stimulated with either PMA plus ionomycin, an unrelated peptide, or the N specific peptide. D. Intracellular accumulation of IFN-γ and IFN-γ, IL-2 and TNF-α in both CD8^+^ T and CD4^+^ T cells from cells isolated from BALs of mice injected with vectors expressing either Nef^mut^ or Nef^mut^/S1. Cells isolated from BALs of five injected mice were pooled, and incubated o.n. with either the S1 peptide pool, or an unrelated pool at the final concentration of 1 μg/ml for each peptide in the presence of brefeldin A. Therefore, cells were analyzed by ICS. Shown are percentages of cytokine expressing CD8^+^/CD4^+^ T cells within the CD3^+^ cell populations after subtraction of values measured in cultures treated with the pool of unrelated peptides. The results are representative of two independent experiments.

In conclusion, the immunization with Nef^mut^-based DNA vectors generated a strong antiviral CD8^+^ T cell immune response also in lung airways, i.e., the peripheral tissue most critically involved in the virus-induced respiratory disease.

### Co-injection of DNA vectors expressing diverse SARS-CoV-2 –based fusion proteins results in an additive immunogenicity effect

A potential features of the Nef^mut^-based CTL vaccine platform is the possibility to immunize against different antigens through a single DNA injection. To explore this possibility, mice were injected with combinations of the DNA vectors expressing SARS-CoV-2 antigens fused with Nef^mut^ already proven to be immunogenic. In particular, mice were injected with DNA vectors expressing Nef^mut^ fused with S1 and M, S1 and N, as well as combination of vectors expressing S1, M and N. As control, equal amounts of the DNA vector expressing Nef^mut^ alone were used. Fourteen days after the second injection, cells from both spleens and BALs were isolated and tested for the presence of SARS-CoV-2-specific CD8^+^ T cells.

When in the IFN-γ EliSpot analysis of splenocytes peptides specific for a single antigen were used, immune responses of potency similar to those previously observed in mice injected with single DNA vectors were detected (Fig. 5A). When combinations of specific peptides were added to IFN-γ EliSpot microwells, the resulting CD8^+^ T cell activation appeared consistently higher than that observed using single peptides (Fig. 5A). The highest immune response was detected in splenocytes from triple injected mice tested with peptides specific for the respective SARS-CoV-2 antigens.

**Figure 5.**
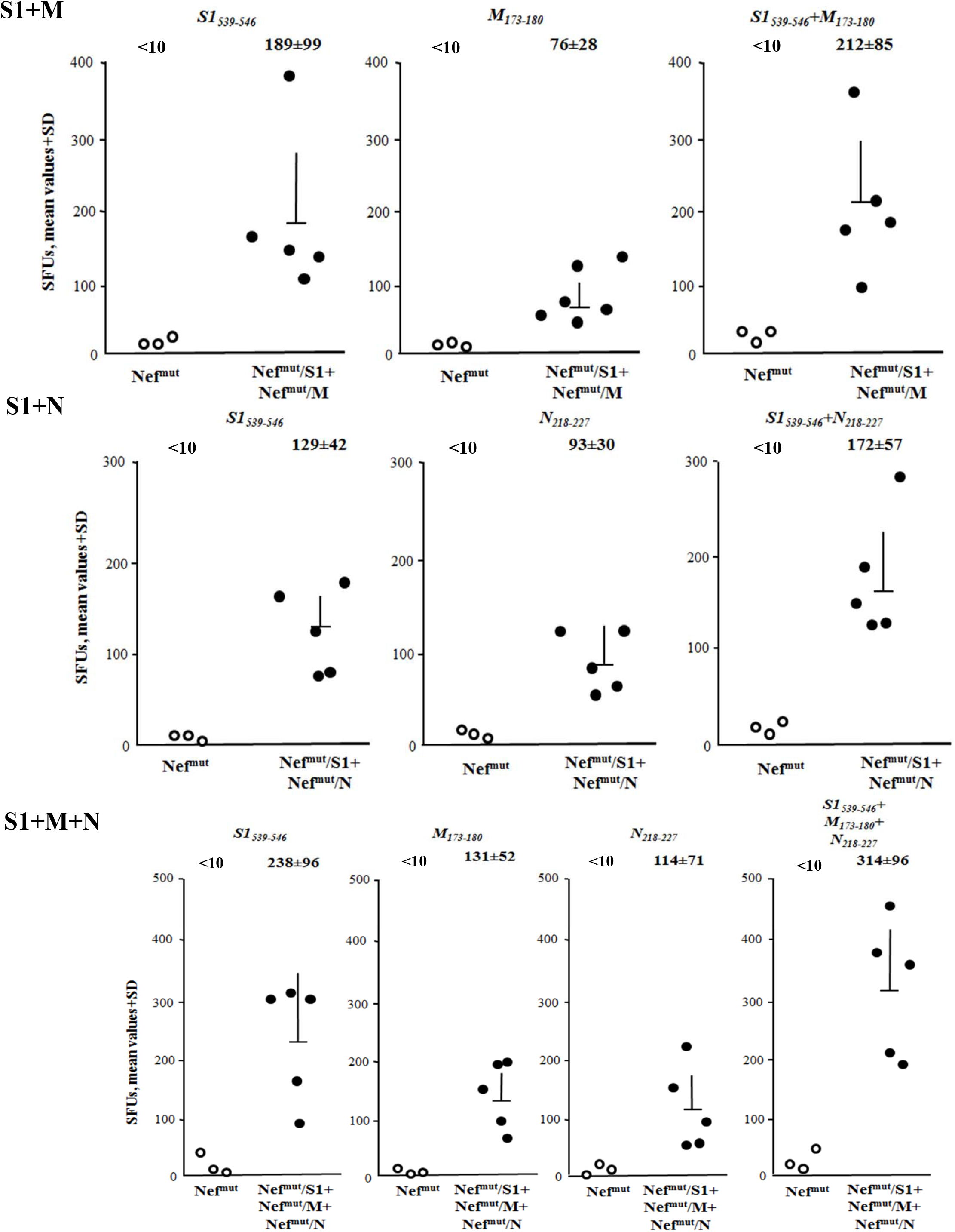
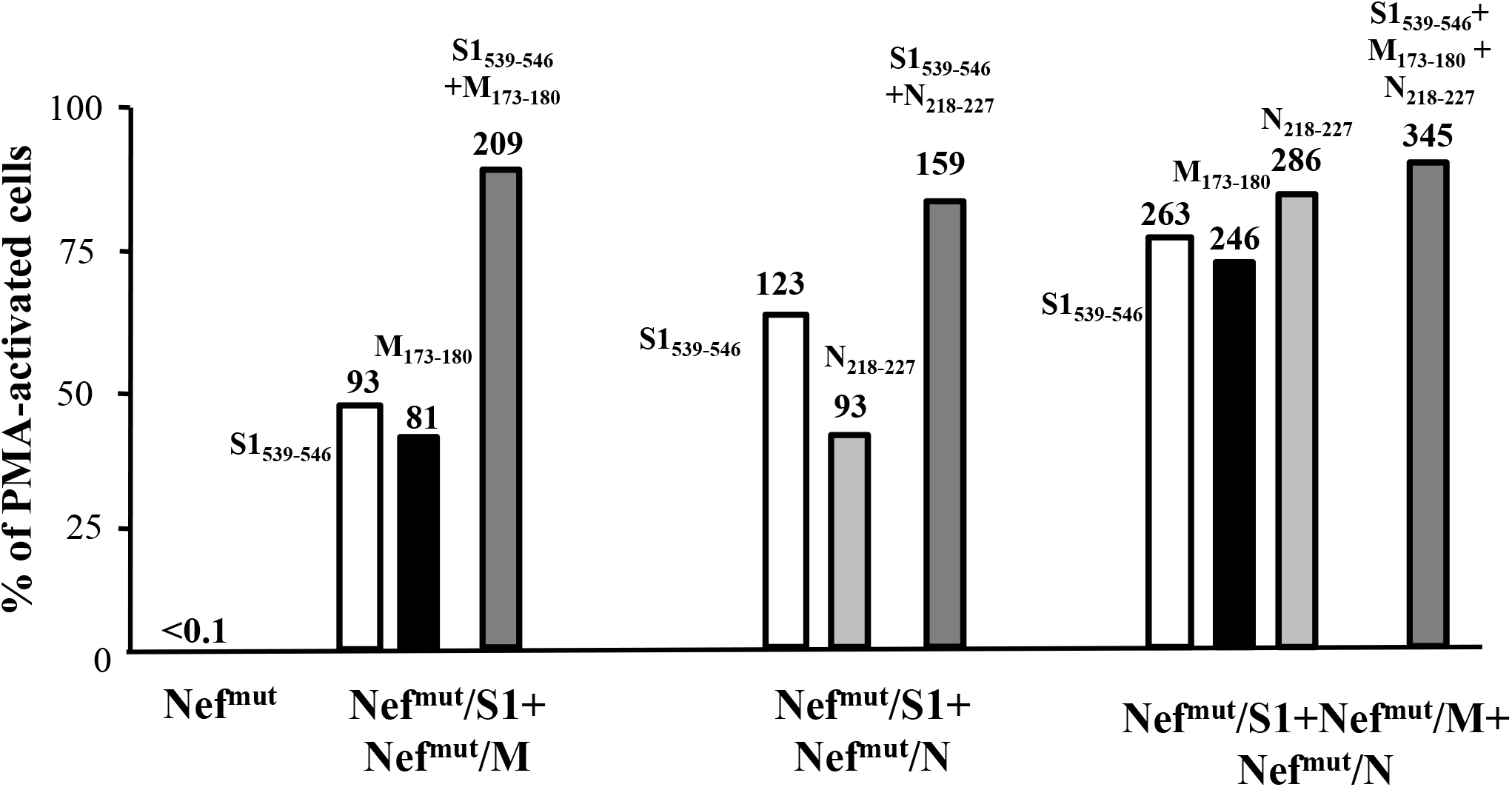
SARS-CoV-2-specific CD8^+^ T cell immune responses induced in both spleens and lungs from mice after i.m. injection of combination of Nef^mut^-based DNA vectors. A. CD8^+^ T cell immune response in either C57 Bl/6 or, for S2 immunization only, Balb/c mice inoculated i.m. with either the DNA vector expressing Nef^mut^ alone (3 mice) or the indicated combinations of vectors expressing Nef^mut^ fused with the SARS-CoV-2 antigens (5 mice per group). The CD8^+^cell immune responses were measured using the indicated peptides or combination thereof. At the time of sacrifice, 2.5×10^5^ splenocytes were incubated o.n. with or without 5 μg/ml of either unrelated or SARS-CoV-2-specific peptides in triplicate IFN-γ EliSpot microwells. Shown are the numbers of SFU/well calculated as mean values of triplicates after subtraction of mean values measured in wells of splenocytes treated with unspecific peptides. Reported are intragroup mean values + standard deviations also. B. Evaluation of the percentages of SFUs detected in IFN-γ EliSpot microwells seeded with 10^5^ cells from BALs pooled from mice injected with the indicated combinations of DNA vectors in the presence of virus-specific peptides compared to PMA plus ionomycin treatment. Conditions comprising either single peptides or combinations thereof were tested. Samples tested in the presence of unrelated peptides scored at background levels. Absolute SFU numbers are also indicated. The results are representative of two independent experiments.

Consistently, the SARS-CoV-2-specific CD8^+^ T cell immune response detected in cells from BALs proved to be quite high in mice injected with DNA combinations (Fig. 5B). Also in this case, the strongest response was detected in cells from mice co- injected with S1-, M- and N-expressing vectors.

The presence of high levels of virus-specific CD8^+^ T cells in lungs represents a strong value added for the here proposed anti-SARS-CoV-2 vaccine strategy. More in general, these data open the way towards the development of CD8^+^ T cell vaccination strategies against multiple antigens of the same and, theoretically, other pathogens also.

### SARS-CoV-2 N specific CTLs are generated upon challenge of human DCs with engineered EVs

The feasibility of the Nef^mut^-based vaccine strategy relies on the possibility to induce virus-specific CTLs in human cells. It was tested by cross-priming experiments carried out by co-cultivating human PBLs with autologous DCs previously challenged with EVs engineered with a SARS-CoV-2 antigen. In detail, human iDCs (Fig. 6A) were challenged with similar amounts (as evaluated by the Alix signal detected in western blot analysis) (Fig. 6B) of EVs uploading either Nef^mut^ or Nef^mut^/N. After overnight culture in the presence of LPS, DCs were put in co-culture with autologous PBLs for a week. Afterwards, the stimulation was repeated and, after an additional week, the presence of SARS-CoV-2-specific CTLs was tested in two ways, i.e., through CD107a and trogocytosis assays.

**Figure 6.**
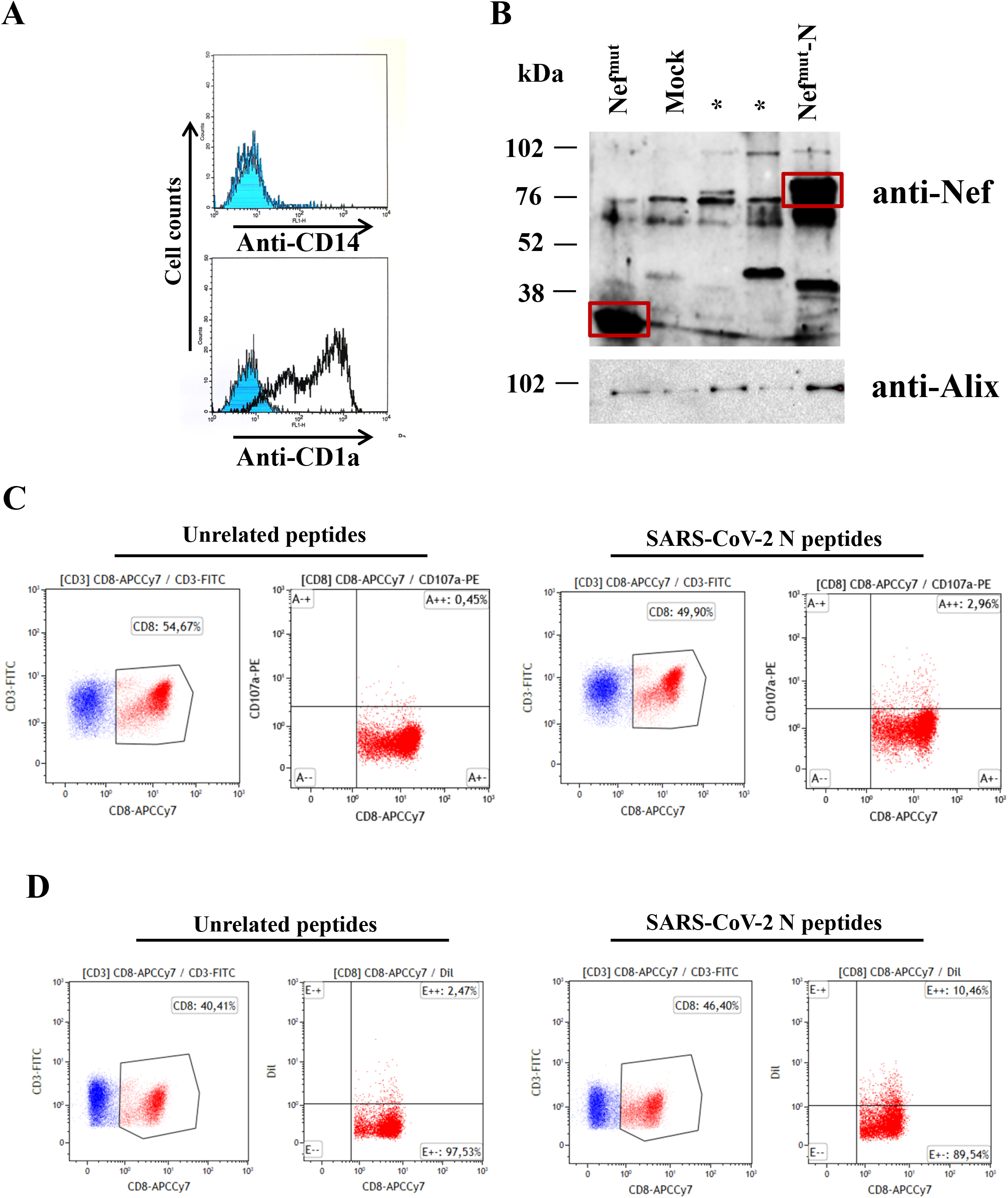
CTL activity as detected in CD8^+^ T lymphocytes co-cultivated with DCs challenged with engineered EVs. A. Phenotype analysis of PBMC-derived monocytes after 5 days of cultivation in the presence of both GM-CSF and IL-4. Cells were tested for the expression of both CD14 (i.e., marker of monocyte-macrophages), and the DC-related CD1a marker. The data are representative of four independent experiments. B. Western blot analysis of equal volumes of buffer where purified EVs were resuspended after differential centrifugations of the supernatants from HEK293T cells transfected 48 hours before with the DNA vector expressing Nef^mut^/N. As control, conditions from mock-transfected cells as well as cells transfected with Nef^mut^ alone were included. Polyclonal anti-Nef Abs served to detect Nef^mut^-based products, while Alix were revealed as markers for EVs. Relevant signals are highlighted. Molecular markers are given in kDa. The results are representative of two independent experiments. Lanes marked with * indicate samples not relevant for the experiments with human cells. C. CD107a FACS analysis on HLA-A.02 PBLs recovered after cross-priming with DCs challenged with Nef^mut^/N engineered EVs. PBLs were analyzed 5 hours after co-culture with HLA-A.02 MCF-7 cells pre-treated with either unrelated of SARS-CoV-2-specific peptides. The results are representative of two independent experiments carried out with cells from different HLA-healthy donors. Percentages of both CD8^+^ and CD107a^+^/CD8^+^ cells are indicated. Percentages of CD8^+^/CD107a^+^ double positive cells measured in PBLs previously co-cultivated with DC challenged with Nef^mut^-engineered EVs remained at background levels. Gating strategy: co-cultures labeled with both FITC-conjugated anti-CD3 and APCCy7-conjugated anti-CD8 mAbs were selected for CD3 expression (for PBL identification), and then the CD8^+^ cell sub-populations were identified within CD3^+^ cells. Quadrants were set on the basis of the fluorescence detected after cell labeling with relevant IgG isotypes. D. Trogocytosis analysis carried out with CD8^+^ T lymphocytes recovered after cross-priming co-cultures. FACS analysis of PBLs recovered as described for panel C, and co-cultivated with CM-Dil-labeled MCF-7 cells. Gating strategy: co-cultures labeled with both FITC-conjugated anti-CD3 and APCCy7-conjugated anti-CD8 mAbs were selected for CD3 expression (for PBL identification), and then the CD8^+^ cells were identified within CD3^+^ cell populations. CD8-labeled cells were finally scored for the CM-Dil-specific fluorescence. Quadrants were set on the basis of the background fluorescence as detected after cell labeling with relevant IgG isotypes. Data are representative of two independent experiments carried out with cells from different HLA-A.02 healthy donors.

The virus-specific cytolytic activity of CD8^+^ T lymphocytes was first assessed through FACS analysis of CD107a/LAMP-1, i.e., a well characterized cell membrane marker of CTL degranulation (*41*). To this end, lymphocytes recovered after cross-priming cultures were co-cultivated for five hours with syngeneic cells (i.e., MCF-7) previously treated with either unrelated or N-specific peptides. We noticed an increase of CD107a expression within the CD8^+^ T cells from PBLs co-cultivated with DCs challenged with N-engineered EVs compared to those co-cultivated with DCs incubated with control EVs (Fig. 6C).

Antigen-specific CTLs can capture plasma membrane fragments from target cells while exerting the cytotoxic activity through a phenomenon referred to as trogocytosis (*42*). The membrane capture is T-cell receptor dependent (*42–45*), epitope specific (*45, 46*), and is exerted by lymphocyte clones endowed with the highest cytotoxic functions (*47*).

At the end of cross-priming cultures, PBLs were put in co-culture with peptide-treated MCF-7 cells previously labeled with CM-Dil, i.e., a red-fluorescent molecule specifically intercalating within cell membrane bilayers. After 5 hours of incubation, the co-cultures were analyzed by FACS for the presence of CD8^+^/Dil^+^ T lymphocytes. As represented in fig. 6D, and consistently with data obtained with the CD107a assay, in the presence of N-specific peptides a significant higher percentage of red-fluorescent cells was reproducibly observed within the CD8^+^ T cells from PBLs co-cultivated with DCs challenged with N-engineered EVs compared to those co-cultivated with DCs incubated with control EVs.

Through these data, we gained the proof-of-principle that EVs engineered with a SARS-CoV-2 antigen have the potential to elicit a virus-specific CTL immune response in humans. This finding is of obvious relevance in view of a possible translation into the clinic of the Nef^mut^-based anti-SARS-CoV-2 vaccine strategy.

## Discussion

Both experimental and clinical evidences already demonstrated the key role of CTLs in the mechanism of protection induced by a number of vaccine preparations against acutely infecting viruses (*48–64*). For instance, in non-human primates the vaccine-induced immunity against Ebola virus is marked by the protective effect largely mediated by CD8^+^ T-cells (*65*). Similarly, a key role of CD8^+^ T-cells in vaccine-induced protection against West Nile Virus has been demonstrated (*66*). Also in the case of infections of human Coronaviruses, CD8^+^ T-cell immunity plays a relevant role in the recovery from SARS-CoV, MERS-CoV (*48*), and SARS-CoV-2 (*19*) infections.

The majority of current efforts towards the development of SARS-CoV-2 vaccines are focused on the induction of antibodies against the S protein. Although the correlates of protection against SARS-CoV-2 are still unknown, results from studies on the related SARS-CoV indicated that the virus-specific antibody response was short lived. In fact, in SARS-CoV-patients, virus-specific IgM and IgA lasted less than 6 months, while virus-specific IgG titers peaked 4 months post-infection and markedly declined after 1 year at best (*5–11*). The risk of losing this protection over time was further indicated in a 6-year follow-up study showing the absence of peripheral memory B cell responses in recovered SARS patients (*12*).

Another serious potential obstacle for SARS-CoV-2 vaccine development is the risk of triggering antibody-dependent enhancement (ADE) of virus infection and/or immunopathology considering that vaccine-induced ADE has been documented in the case of SARS-CoV infections (*67–69*), and just recently suggested for SARS-CoV-2 (*70*). Thus, without a full understanding of the mechanisms underlying protective immunity, many fear that some vaccines might worsen the disease rather than prevent it, echoing the disastrous effects of the Dengvaxia tetravalent yellow fever-dengue antibody-generating vaccine (*71*).

Following a prime-boost immunization approach, Channappanavar and coll. showed that CD8^+^ T-cells specific for a single SARS-CoV immunodominant epitope protected mice from an otherwise lethal dose of virus in the absence of neutralizing antibodies (*72*). The same study demonstrated the lack of memory CD8^+^ T-cell-mediated immunopathology, suggesting that the induction of these cells was safe (*72*). On the other hand, unlike patient waning serum antibody levels, CD8^+^ T-cell responses against S and N proteins can still be detected in peripheral blood of recovered SARS-CoV patients even 11 years post-infection (*12–17*). Recently, Grifoni and coll. demonstrated robust T cell responses against S, M, and N proteins in 20 COVID-19 convalescent patients (*18*). Circulating SARS-CoV-2-specific CD8^+^ and CD4^+^ T cells were found in 70% and 100% of patients, respectively (*18*). Another study identified the most part of SARS-CoV-2-specific epitopes recognized by memory CD8^+^ T cells from COVID-19 patients in both N and ORF1ab proteins (*19*). Taken together, these evidences suggest that a strong memory CD8^+^ T-cell response should be a component of any vaccine regimen for human CoVs, possibly in combination with immunogens inducing safe neutralizing antibodies.

As from previous immunogenicity studies with DNA vectors expressing Nef^mut^ fused with several viral and tumor antigens, HPV-16 E7 reproducibly appeared the antigen eliciting the most potent CD8^+^ T cell response (*29, 35*). Here, we provide evidence that DNA vectors expressing single SARS-CoV-2 antigens elicited a CD8^+^ T cell immune response at least comparable to that induced by Nef^mut^/E7 (*36*). Most striking, valuable levels of virus-specific CD8^+^ T cells were identified in cells from BALs of immunized mice. Resident CD8^+^ T cells in lung are basically maintained independently of the pool of circulating CD8^+^ T cells, undergoing homeostatic proliferation to replenish the continuous loss of cells through intraepithelial migration toward lung airways. In view of the quite high percentages of virus-specific CD8^+^ T cells we detected in BALs, it is conceivable that these cells originated by activation occurred at local, e.g., mediastinal, lymph nodes rather than by diffusion of circulating virus-specific CD8^+^ T cells. This hypothesis is consistent with the biological properties of immunogenic EVs, which are expected to freely circulate into the body thereby targeting, as already documented (*73*), spleen, liver, and lung. Upon EV capture, tissue-resident APCs would migrate to local lymph nodes thereby switching the processes leading to antigen-specific CD8^+^ T cell activation. Even if the underlying mechanism needs to be elucidated, the results we obtained with BALs should be considered of overwhelming relevance for a cell-mediated vaccine strategy conceived to battle a respiratory virus.

The immune response we observed by co-injecting different DNA vectors appeared reproducibly higher than that obtained by injecting each DNA vector alone. No negative interferences among the diverse antigens were observed by testing immune responses with cells isolated from spleens and, most important, BALs. Hence, the Nef^mut^-based vaccine platform offers the option to elicit a simultaneous CD8^+^ T cell response against different antigens, which were proven to be effective in peripheral tissues as well. The results we obtained with *ex vivo* human cells represent the proof-of-principle that the anti-SARS-CoV-2 vaccine strategy based on engineered EVs could be readily translated into the clinic.

The overall significance of our findings would greatly benefit from data of virus challenge, for instance, on hACE-transgenic mice immunized by Nef^mut^-based DNA vectors. Notwithstanding, a number of original achievements has been already obtained through our studies, in particular: i) strong CD8^+^ T immunity was generated by DNA vectors expressing either SARS-CoV-2 S1, S2, M or N proteins fused with Nef^mut^; ii) this immune responses were detectable in lung airways also, and iii) the CD8^+^ T immune response can be elicited against different antigens through a single injection. All these findings would have significance also whether applied to other infectious or tumor antigens.

Taken together DNA vectors expressing the products of fusion between Nef^mut^ and SARS-CoV-2 antigens can be considered candidates for new vaccine strategies aimed at inducing anti-SARS-CoV-2 CTLs.

## Materials and Methods

### DNA vector synthesis

ORFs coding for Nef^mut^ fused with S1, S2, M, and N SARS-CoV-2 proteins were cloned into pVAX1 plasmid (Thermo Fisher), i.e., a vector approved by FDA for use in humans. To obtain the pVAX1 vector expressing Nef^mut^, its ORF was cloned into *Nhe* I and *EcoR* I sites (Fig. 1A). To recover vectors expressing Nef^mut^-based fusion products, an intermediate vector referred to as pVAX1/Nef^mut^fusion was produced (Fig. 1A). Here, the whole Nef^mut^ ORF deprived of its stop codon was followed by a sequence coding a GPGP linker including a unique *Apa* I restriction site. In this way, sequences comprising the *Apa* I site at their 5’ end and the *Pme* I one at their 3’ end were fused in frame with Nef^mut^ ORF (Fig. 1). Stop codons of SARS-CoV-2-related sequences were preceded by sequences coding for a DYKDDDK epitope tag (flag-tag). SARS-CoV-2 sequences were optimized for expression in human cells through the GeneSmart software from Genescript. All these vectors were synthesized by Explora Biotech. The pTargeT vector expressing the Nef^mut^/HPV16-E7 fusion protein was already described (*74*).

### Cell cultures and transfection

Human embryonic kidney (HEK) 293T cells (ATCC, CRL-11268) were grown in DMEM (Gibco) plus 10% heat-inactivated fetal calf serum (FCS, Gibco). Transfection assays were performed using Lipofectamine 2000 (Invitrogen, Thermo Fisher Scientific)-based method.

### EV isolation

Cells transfected with vectors expressing the Nef^mut^-based fusion proteins were washed 24 h later, and reseeded in medium supplemented with EV-deprived FCS. The supernatants were harvested from 48 to 72 h after transfection. EVs were recovered through differential centrifugations (*75*) by centrifuging supernatants at 500×*g* for 10 min, and then at 10,000×*g* for 30 min. Supernatants were harvested, filtered with 0.22 μm pore size filters, and ultracentrifuged at 70,000×*g* for l h. Pelleted vesicles were resuspended in 1×PBS, and ultracentrifuged again at 70,000×*g* for 1 h. Afterwards, pellets containing EVs were resuspended in 1:100 of the initial volume.

### Western blot analysis

Western blot analyses of both cell lysates and EVs were carried out after resolving samples in 10% sodium dodecyl sulfate-polyacrylamide gel electrophoresis (SDS-PAGE). In brief, the analysis on cell lysates was performed by washing cells twice with 1×PBS (pH 7.4) and lysing them with 1× SDS-PAGE sample buffer. Samples were resolved by SDS-PAGE and transferred by electroblotting on a 0.45 μM pore size nitrocellulose membrane (Amersham) overnight using a Bio-Rad Trans-Blot. EVs were lysed and analyzed as described for cell lysates. For immunoassays, membranes were blocked with 5% non-fat dry milk in PBS containing 0.1% Triton X-100 for 1 h at room temperature, then incubated overnight at 4 °C with specific antibodies diluted in PBS containing 0.1% Triton X-100. Filters were revealed using 1:1,000-diluted sheep anti-Nef antiserum ARP 444 (MHRC, London, UK), 1:500-diluted anti-β-actin AC-74 mAb from Sigma, 1:500 diluted anti-Alix H-270 polyclonal Abs from Santa Cruz, and 1:1,000 diluted anti-flag M2 mAb from Sigma.

### Mice immunization

Both 6-weeks old C57 Bl/6 and, for S2 immunization (in view of the lack of already characterized H2^b^ immunodominant S2 epitopes), Balb/c female mice were obtained from Charles River. They were housed at the Central Animal Facility of the Istituto Superiore di Sanità, as approved by the Italian Ministry of Health, authorization n. 565/2020 released on June, 3, 2020. The day before the first inoculation, microchips from DATAMARS were inserted sub cute at the back of the neck between the shoulder blades on the dorsal midline. DNA vector preparations were diluted in 30 μL of sterile 0.9% saline solution. Both quality and quantity of the DNA preparations were checked by 260/280 nm absorbance and electrophoresis assays. Mice were anesthetized with isoflurane as prescribed in the Ministry authorization. Each inoculum volume was therefore measured by micropipette, loaded singly into a 1 mL syringe without dead volume, and injected into mouse right quadriceps. Immediately after inoculation, mice underwent electroporation at the site of injection through the Agilpulse BTX device using a 4-needle array 4 mm gap, 5 mm needle length, with the following parameters: 1 pulse of 450 V for 50 μsec; 0.2 msec interval; 1 pulse of 450 V for 50 μsec; 50 msec interval; 8 pulses of 110 V for 10 msec with 20 msec intervals. The same procedure was repeated for the left quadriceps of each mouse. Immunizations were repeated after 14 days. Fourteen days after the second immunization, mice were sacrificed by either cervical dislocation or CO_2_ inhalation, in both cases following the recommendations by the Ministry authorization protocol.

### Cell isolation from immunized mice

Spleens were explanted by qualified personnel of the ISS Central Animal Facility, and placed into a 2 mL Eppendorf tubes filled with 1 mL of RPMI 1640 (Gibco), 50 μM 2-mercaptoethanol (Sigma). Spleens were transferred into a 60 mm Petri dish containing 2 mL of RPMI 1640 (Gibco), 50 μM 2-mercaptoethanol (Sigma). Splenocytes were extracted by incising the spleen with sterile scissors and pressing the cells out of the spleen sac with the plunger seal of a 1 mL syringe. After addition of 2 mL of RPMI medium, cells were transferred into a 15 mL conical tube, and the Petri plate was washed with 4 mL of medium to collect the remaining cells. After a three-minute sedimentation, splenocytes were transferred to a new sterile tube to remove cell/tissue debris. Counts of live cells were carried out by the trypan blue exclusion method. Fresh splenocytes was resuspended in RPMI complete medium, containing 50 μM 2-mercaptoethanol and 10% FBS, and tested by IFN-γ EliSpot assay.

For bronchoalveolar lavages, mice were sacrificed by CO_2_ inhalation, placed on their back, and dampened with 70% ethanol. Neck skin was opened to the muscles by scissors, and muscles around the neck and salivary glands were gently pulled aside with forceps to expose the trachea. A 15 cm long surgical thread was then placed around the trachea and a small hole was cut by fine point scissors between two cartilage rings. A 22 G×1” Exel Safelet Catheter was carefully inserted about 0.5 cm into the hole, and then the catheter and the trachea were firmly tied together with the suture. A 1 mL syringe was loaded with 1 mL of cold PBS and connected to the catheter. The buffered solution was gently injected into the lung and aspirated while massaging mouse torax. Lavage fluid was tranferred to a 15 mL conical tube on ice, and lavage repeated two more times (*76, 77*). Total lavage volume was approximately 2.5 mL/mouse. Cells were recovered by centrifugation, resuspeded in cell culture medium, and counted.

### IFN-γ EliSpot analysis

A total of 2.5×10^5^ live cells were seeded in each IFN-γ EliSpot microwell. Cultures were run in triplicate in EliSpot multiwell plates (Millipore, cat n. MSPS4510) pre-coated with the AN18 mAb against mouse IFN-γ (Mabtech) in RPMI 1640 (Gibco), 10% FCS, 50 μM 2-mercaptoethanol (Sigma) for 16 h in the presence of 5 μg/mL of the following CD8-specific peptides: HPV-16 E7 (H2-K^b^): 21-28 DLYCYEQL (*78*); 49-57 RAHYNIVTF (*78*); 67-75 LCVQSTHVD (*79*). SARS-CoV-2 S1 (H2-K^b^): 525-531 VNFNFNGL (*80*); SARS-CoV-2 S2 (H2-K^d^): 1079-1089 PAICHDGKAH (*81*); SARS-CoV-2 M (H2-K^b^): 173-180 RTLSYYKL (*82*); SARS-CoV-2 N (H2-K^b^): 219-228 ALALLLLDRL (*82*). As negative controls, 5 μg/mL of either H2-K^b^ or H2-K^d^-binding peptides were used. More than 70% pure preparations of the peptides were obtained from Elabioscences. SARS-CoV-2 peptide collections were obtained from BEI resources. Peptide pools were adjusted at the concentration of 50 μg/mL for each peptide, and used at the final concentration of 1 μg/mL each. For cell activation control, cultures were treated with 10 ng/mL PMA (Sigma) plus 500 ng/mL of ionomycin (Sigma). PeptTivator NS3 (Miltenyi) (i.e., a pool of HCV genotype 1b NS3 peptides consisting in 15-mers with 11 aa overlap covering the whole NS3 sequence) was used as negative control. After 16 hours. cultures were removed, and wells incubated with 100 μL of 1 μg/ml of the R4-6A2 biotinylated anti-IFN-γ (Mabtech) for 2 hours at r.t. Wells were then washed and treated for 1 hour at r.t. with 1:1000 diluted streptavidine-ALP preparations from Mabtech. After washing, spots were developed by adding 100 μL/well of SigmaFast BCIP/NBT, Cat. N. B5655. The spot-forming cells were finally analyzed and counted using an AELVIS EliSpot reader.

### Intracellular staining

Cells from BALs were seeded (2×10^6^/mL) in RPMI medium, 10% FCS, 50 μM 2-mercaptoethanol (Sigma), and 1 μg/mL Brefeldin A (BD Biosciences). Control conditions were carried out either by adding 10 ng/ml PMA (Sigma) and 1 μg/mL ionomycin (Sigma), or with unrelated peptides. After 16 hours, cultures were stained with 1 μl of LIVE/DEAD Fixable Aqua Dead Cell reagent (Invitrogen ThermoFisher) in 1 mL of PBS for 30 minutes at 4 °C and washed twice with 500 μl of PBS. To minimize nonspecific staining, cells were pre-incubated with 0.5 μg of Fc blocking mAbs (i.e., anti-CD16/CD32 antibodies, Invitrogen/eBioscience) in 100 μL of PBS with 2% FCS for 15 minutes at 4 °C. For the detection of cell surface markers, cells were stained with 2 μL of the following Abs: FITC conjugated anti-mouse CD3, APC-Cy7 conjugated anti-mouse CD8a, and PerCP conjugated anti-mouse CD4 (BD Biosciences) and incubated for 1 hour at 4 °C. After washing, cells were permeabilized and fixed through the Cytofix/Cytoperm kit (BD Biosciences) as per the manufacturer’s recommendations, and stained for 1 hour at 4 °C with 2 μl of the following Abs: PE-Cy7 conjugated anti-mouse IFN-γ, PE conjugated anti-mouse IL-2 (Invitrogen eBioscience), and BV421 rat anti-mouse TNF-α BD Biosciences in a total of 100 μL of 1× Perm/Wash Buffer (BD Biosciences). After two washes, cells were fixed in 200 μL of 1× PBS/formaldehyde (2% v/v). Samples were then assessed by a Gallios flow cytometer and analyzed using Kaluza software (Beckman Coulter).

Gating strategy was as follows: live cells as detected by Aqua LIVE/DEAD dye vs. FSC-A, singlet cells from FSC-A vs. FSC-H (singlet 1) and SSC-A vs SSC-W (singlet 2), CD3 positive cells from CD3 (FITC) vs. SSC-A, CD8 or CD4 positive cells from CD8 (APC-Cy7) vs. CD4 (PerCP). The CD8^+^ cell population was gated against APC-Cy7, PE, and BV421 to observe changes in IFN-γ, IL-2, and TNF-α production, respectively. Boolean gates were created in order to determine any cytokine co-expression pattern.

### Cross-priming assay

Monocytes were isolated from HLA-A.02 peripheral blood mononuclear cells (PBMCs) by immune magnetic protocol. Human immature dendritic cells (iDCs) were obtained after 5 to7 days of culture of monocytes in the presence of both IL-4 and GM-CSF. Immature DCs were challenged by engineered EVs uploading either Nef^mut^/N or Nef^mut^ alone isolated from supernatants of HEK293T transfected cells. After an overnight incubation, iDCs were matured by LPS treatment for 24 hours. Thereafter, DCs were washed, and co-cultured with autologous peripheral blood lymphocytes (PBLs) in a 1:10 cell ratio. A week later, the stimulation procedure was repeated, and, after an additional week, CD8^+^ T cells were recovered for downstream assays.

### Activation-induced degranulation and trogocytosis assays

Activation-induced degranulation was measured by evaluating CD107a surface expression as described (*41*). Briefly, 2×10^5^ PBLs recovered from cross-primed cultures were co-cultivated for 5 hours with equal number of HLA-matched target cells, i.e., MCF-7 previously treated with 1 μg/mL of either SARS-CoV-2 N-specific or unrelated HLA-A.02 binding peptides, in the presence of both PE-conjugated anti-CD107a (BD-Pharmingen) and 0.7 μg/mL monensin (GolgiStop, BD). The gated CD8^+^ T cell populations were then analyzed by FACS for the detection of CD107a-related fluorescence.

For trogocytosis assay, MCF-7 cells were labeled with the fluorescent lipophilic dye CM-Dil (Molecular Probes) according to the manufacturer's instructions with minor modifications. In detail, 10^6^ target cells were resuspended in 1 mL of 1×PBS in the presence of 1 μM CM-Dil, and incubated for 5 min at 37 °C, following by an additional incubation of 15 min at 4 °C. Cold RPMI medium containing 10% AB human serum was added to stop labeling (1:1 volume) for 10 min at 4 °C. Then, cells were washed three times with 1×PBS containing 1% AB human serum. Following resuspension in complete medium, labeled target cells were cultured for 4 hours in the presence of 1 μg/mL of either SARS-CoV-2 N-specific or unrelated HLA-A.02 binding peptides, and finally co-cultured with primed PBLs (5×10^5^ per well in 200 μL of total volume) in U-shaped 96-well plates in a 1:5 ratio. After incubation of 5 hours at 37 °C, cells were washed twice in 1×PBS containing 0.5 mM EDTA to ensure cell dissociation, resuspended in 1×PBS supplemented with 0.5% bovine serum albumin, and stained with appropriate Abs for FACS analysis. Dead cells were stained by adding Aqua LIVE/DEAD dye. HLA-A.02-binding SARS-CoV-2 N-specific peptides were: 159-167 LQLPQGTTL, 219-227 ALLLLDRL, 316-324 GMSRIGMEV, and 351-359 ILLNKHIDA (*83*).

### Statistical analysis

When appropriate, data are presented as mean + standard deviation (SD). In some instances, the Mann-Whitney U test was used. *p*<0.05 was considered significant.

## Ackowledgements

The following reagents were obtained through BEI Resources, NIAID, NIH: Peptide Array, SARS-Related Coronavirus 2 Spike (S) Glycoprotein, NR-52402, Nucleocapside (N) Protein, NR-52404, and Membrane (M) Protein, NR-52403. We thank Andrea Giovannelli, Antonio Di Virgilio, and Pietro Arciero, Istituto Superiore di Sanità, for technical support.

